# Microstructure-Based Modeling of Primary Cilia Mechanics

**DOI:** 10.1101/2023.07.14.549117

**Authors:** Nima Mostafazadeh, Andrew Resnick, Y.-N. Young, Zhangli Peng

## Abstract

A primary cilium, made of nine microtubule doublets enclosed in a cilium membrane, is a mechanosensing organelle that bends under an external mechanical load and sends an intracellular signal through transmembrane proteins activated by cilium bending. The nine microtubule doublets are the main load-bearing structural component, while the transmembrane proteins on the cilium membrane are the main sensing component. No distinction was made between these two components in all existing models, where the stress calculated from the structural component (nine microtubule doublets) was used to explain the sensing location, which may be totally misleading. For the first time, we developed a microstructure-based primary cilium model by considering these two components separately. First, we refined the analytical solution of bending an orthotropic cylindrical shell for individual microtubule, and obtained excellent agreement between finite element simulations and the theoretical predictions of a microtubule bending as a validation of the structural component in the model. Second, by integrating the cilium membrane with nine microtubule doublets, we found that the microtubule doublets may twist significantly as the whole cilium bends. Third, besides being cilium-length-dependent, we found the mechanical properties of the cilium are also highly deformation-dependent. More important, we found that the cilium membrane near the base is not under pure in-plane tension or compression as previously thought, but has significant local bending stress. This challenges the traditional model of cilium mechanosensing, indicating that transmembrane proteins may be activated more by membrane curvature than membrane stretching. Finally, we incorporated imaging data of primary cilia into our microstructure-based cilium model, and found that comparing to the ideal model with uniform microtubule length, the imaging-informed model shows the nine microtubule doublets interact more evenly with the cilium membrane, and their contact locations can cause even higher bending curvature in the cilium membrane than near the base.

**SIGNIFICANCE:** Factors regulating the mechanical response of a primary cilium to fluid flow remain unclear. Modeling the microtubule doublet as a composite of two orthotropic shells and the ciliary axoneme as an elastic shell enclosing nine such microtubule doublets, we found that the length distribution of microtubule doublets (inferred from cryogenic electron tomography images) is the primary determining factor in the bending stiffness of primary cilia, rather than just the ciliary length. This implies ciliary-associated transmembrane proteins may be activated by membrane curvature changes rather than just membrane stretching. These insights challenge the traditional view of ciliary mechanosensation and expands our understanding of the different ways in which cells perceive and respond to mechanical stimuli.

## 1 INTRODUCTION

The primary cilium is a membrane-encased microtubule-based cellular organelle that protrudes out of the apical side of the cell when the mother centriole (microtubule organization center, MTOC) docks to the plasma membrane (1). As the cilium grows, the axoneme consisting of nine microtubule doublets lengthen while enclosed within a ciliary membrane that is contiguous with the cell membrane yet has its own distinct chemical composition (2). Long speculated to be decorative, the primary cilium is now hypothesized to be a nexus of cellular sensing (3) especially vital to normal functioning in a wide range of mammalian cells. Strong evidence for this hypothesis is the commonly observed cytosolic increase in Calcium when the primary cilium is bent (4). This response was lost with removal of primary cilium (4, 5).

Structural or functional defects in primary cilia (due to genetic disorders (6), for example) may disturb their mechanosensing pathways and are implicated in serious diseases, for example polycystic kidney disease (PKD) (7–12). However, the connection between ciliary-mediated flow responses and overall organismal health and disease is still debated. For example, although a large body of results indicate that the activation of transmembrane calcium channels on cilia are mechanically induced (13–15), the specific molecular (mechanistic) events connecting mechanical bending of a cilium to release of intracellular Calcium remains unclear (16).

Ferreira et al. (17) raised the idea that primary cilia are mechanosensitive organelles due to both ciliary protein composition and ciliary stress distribution under mechanical deflection. Given the difficulties of studying primary cilia function on the molecular scale, numerical modeling could partially relieve the experimental difficulties of understanding the individual and combined mechanical responses from both microtubules and plasma membrane, allowing integration of overall ciliary mechanics and determine local stress distributions for each member of this intricate structure.

The mechanical properties of primary cilia are closely related to those of microtubules, which have been shown to have a length-dependent persistence length (18): Longer microtubules are stiffer than shorter microtubules. Such dependence on the microtubule length can be explained by the anisotropic shell model of a cylindrical shell with a much larger axial modulus than the azimuthal moduli (19). Previously (20) we provided, for the first time, evidence that the persistence length of a ciliary axoneme is also length-dependent; longer cilia are stiffer than shorter cilia, just as microtubules. We demonstrated that this apparent length dependence can be understood by a combination of modeling axonemal microtubules as anisotropic elastic shells and including actomyosin-driven stochastic basal body motion. Our results also demonstrated that it is possible to use observable ciliary dynamics to probe the cytoskeletal dynamics inside the cell.

More importantly, the nine microtubule doublets are the main load-bearing structural component, while the transmembrane proteins on the cilium membrane are the main sensing component. No distinction was made between these two components in all existing models (21–25), where the stress calculated from the structural component (nine microtubule doublets) was used to explain the sensing location, which may be totally misleading, as the stress in bent microtubule doublets may not correlate to the stress in the sensing component (cilium membrane). For the first time, we developed a microstructure-based primary cilium model by considering these two components separately.

In addition, the traditional paradigm of modeling ciliary axoneme assumes that the nine microtubule doublets have the same length as the axonemal centerline (26). However, Sun *et al*. (27) reported a very different micro architecture of epithelial primary cilia, where the nine microtubule doublets, or microtubule complexes (MtCs), have different lengths along the axoneme axis and not all MtCs reach the axoneme tip. Furthermore, some of the MtCs transition to singlets. How would such microstructural architecture of MtCs affect the mechanics of the primary cilium under an external load? Furthermore, how might such architecture of MtCs result in different mechanisms for mechanotransduction of primary cilia? To answer these questions, we combine the improved characterization of cilia with the observed micro structures of MtCs inside the ciliary axoneme to gain deeper understanding of the mechanics of primary cilia and a more complete insight into the underlying mechanisms for their mechanosensing and length regulation.

For this paper, we constructed numerical models for both ciliary plasma membrane and microtubule mechanics using electron-microscopy (EM) image-based architecture from experimental data. We simulated tip-anchored optical tweezer experiments with varying cilium lengths and verified the resulting computed displacements with an analytical continuum shell model solution. We then further analyzed computational results to quantify distinct membrane and axonemal contributions to mechanical responses of primary cilia and identify possible mechanosensing triggers during bending deformation.

## 2 ORTHOTROPIC SHELL THEORY OF MICROTUBULES

Microtubules are tubular structures composed of tubulins (Fig 1a). The basic building block of microtubules is the U-V tubulin heterodimer, which forms after U- and V-tubulin monomers co-crystallize during the nucleation process. These heterodimers then assemble in head-to-tail configuration to produce protofilaments (28). In general, 11 to 15 protofilaments laterally associate to create hollow tube-like structure, and microtubules with 13 protofilaments are the most common (29). Anisotropy in microtubule mechanics arises from the contrast between U-V lateral and longitudinal interactions between tubulins. Although all attachments are non-covalent and mediated through hydrogen and electrostatic bonds (30), binding strength in between tubulins in longitudinal direction is about twice as strong as between adjacent protofilaments (31).

**Figure 1.**
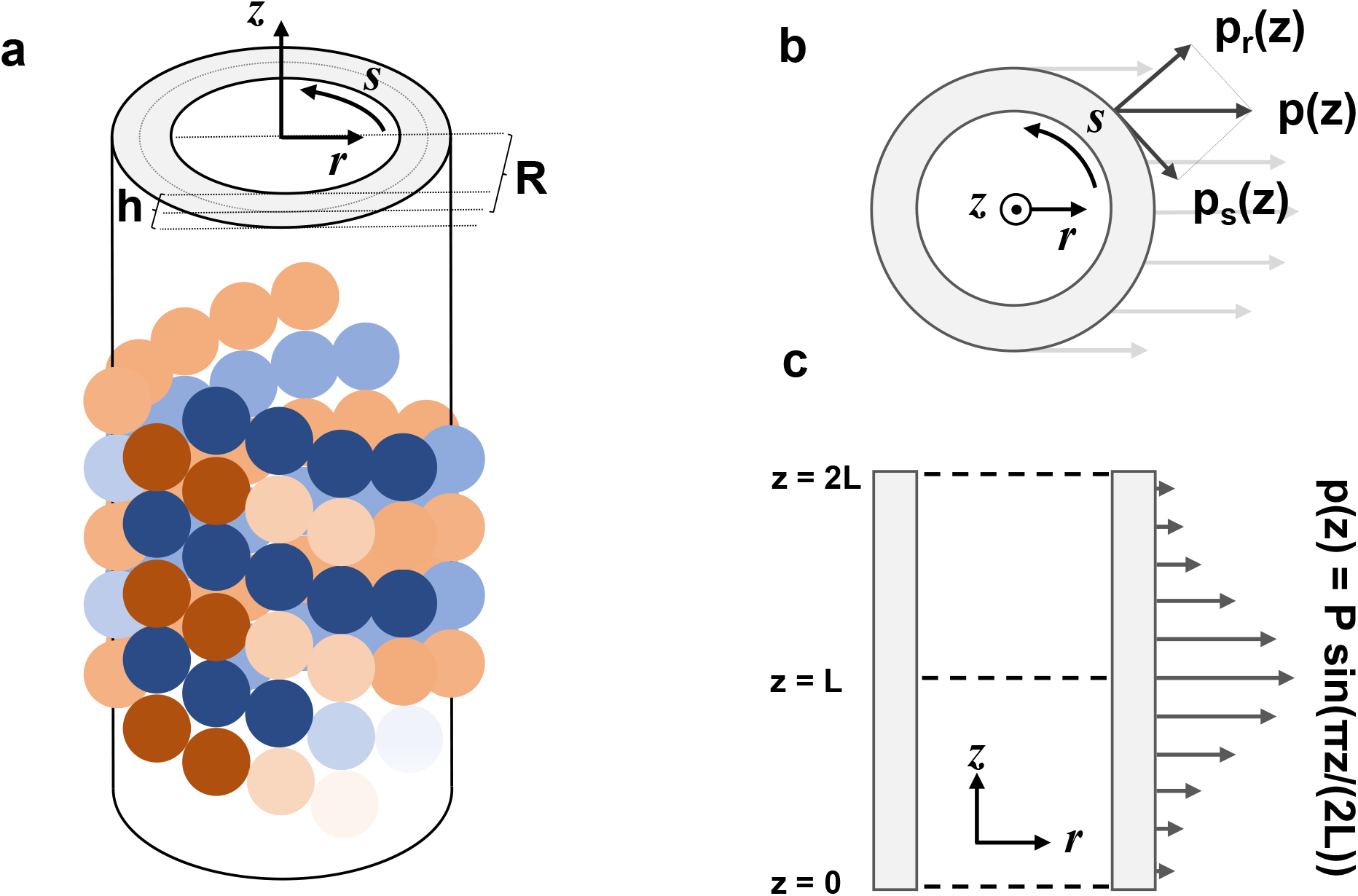
: Illustration of microtubule structure and model geometry. (a): Tubulin proteins are represented as beads. A microtubule is a hollow cylinder of tubulin monomers interconnected through distinct lateral and longitudinal bonds. (b): Schematic of model microtubule shell using cylindrical coordinates of radius *R*, thickness *h* and length *L*. Radius is measured up to middle section in thickness; the inner and outer radii are *R* −*h/*2 and *R* + *h*/2. (c): *p*(*z*), distributed load in height *z* is designated with its radial and circumferential components *p*(*z*)_*r*_ and *p*(*z*)_*s*_. The load is only applied to half of the cylinder surface. (d): Distribution of the total load, *P*, on the *z* axis follows a periodic function for analytical convenience. Our theory is symmetric about the plane z = L, and so by symmetry, the simulated ‘half-problem’ is equivalent to the theory ‘full problem’.

Efforts have been made to characterize elastic properties of microtubules through experiments and simulations. Atomic force microscopy (AFM) indentation (32, 33), optical tweezers (34, 35), and thermal fluctuation wave analysis (36, 37) are employed to estimate the flexural rigidity of microtubule shells. Consequently, Young’s moduli in order of 10^9^ Pa have been reported for certain bending criteria and geometric assumptions. The persistence length of microtubules varies and approaches to a plateau of roughly 5000 ’m as length-to-radius ratio of microtubules increases. For experiments which have conducted at this scale, the longitudinal stiffness of microtubule becomes dominant, on the order of giga Pascals, and can confidently be attributed to longitudinal elastic modulus *E*_*z*_ of microtubules. Weaker bonds in circumferential direction implies smaller *E*_*s*_ compared to *E*_*z*_. Simulations indicate that the magnitude of *E*_*s*_ should be in the range of 1-4 MPa, about 1000 times smaller than *E*_*z*_. Simulations and mathematical analysis were exploited to estimate shear modulus of a shell tube with flexural rigidity of a microtubule, with results varying from 1 kPa to 10 MPa (38, 39). The ranges reported in the literature are presented in Table 1.

**Table 1:** Parameter values for the three components in the numerical model.

| Component | Parameter Symbol | Literature | Used Value |
| --- | --- | --- | --- |
| Microtubule | $R$ | 8 – 12 nm (32) | 10 nm |
| | $h$ | 5 – 6 nm (38) | 5 nm |
| | $E_z$ | 1MPa - 7 GPa (32–37, 40) | 2.6 GPa |
| | $E_s$ | 1 – 4 MPa (38, 39) | 2 MPa |
| | $G$ | 1 kPa - 1 MPa (32, 39) | 7 kPa |
| | $\nu_{zs}$ | 0.3 (38) | 0.3 |
| Lipid bilayer | $B_M$ | $2 \times 10^{-19} J$ (41) | $2 \times 10^{-19} J$ |
| | $K_M$ | $4.5 \times 10^5 \text{ pN}/\mu\text{m}$ (41) | $4.5 \times 10^5 \text{ pN}/\mu\text{m}$ |
| Primary cilium | $R_{CM}$ | $\sim 100 \text{ nm}$ (27) | 100 nm |
| | $r_a$ | n/a | $5/6 R_{CM}$ |
| | $r_b$ | n/a | $4/3 R$ |

Based on the molecular structures of microtubules, we expect the orthotropic shell model to effectively capture the distinct elastic and shear mechanics of microtubule as it bends under a load. For a thin cylindrical shell of length *L*, radius *R* (middle point), and thickness *h*, 3D cylindrical displacements *U, V* and *W* are defined in longitudinal *z*, circumferential *s*, and radial *r* directions, respectively (Fig 1b). In our model formulation, the corresponding strain equations incorporate second-order curvature terms, while all normal and shear radial strains are assumed to be zero, in accordance with the thin shell assumption. Asserting the mid-point radius as the basis (42), the expressions for the strain are

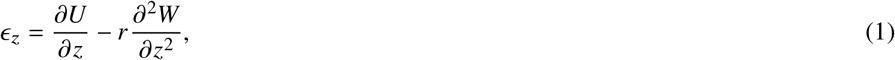

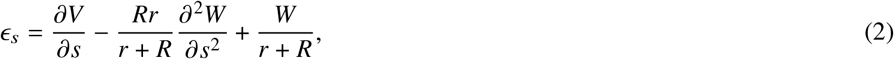

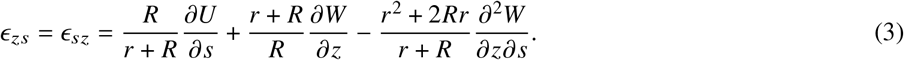

Similarly, the equations for stress are formulated for the longitudinal, circumferential and their shear directions, with the respective elastic moduli *E*_*z*_ and *E*_*s*_, shear modulus *G*, as well as Poisson ratios *v*_*zs*_and *v*_*sz*_(43):

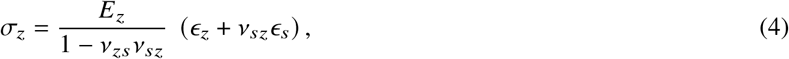

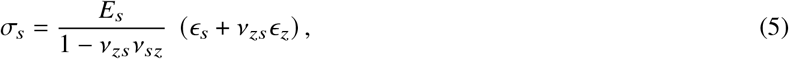

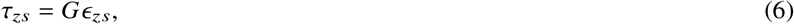

where *E*_*z*_*v*_*sz*_= *E*_*s*_*v*_*zs*_.

Our goal is to obtain an expression of the lateral displacement of an orthotropic shell under a symmetric radial loading *P*, where 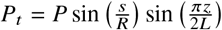 and 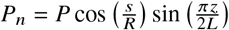 represent the tangential and normal components of the periodically distributed load *P*, respectively (Fig 1c-d). By substituting the stress and strain relations (Eqs. 1-6) into the definitions of moments and internal force, and simplifying the equilibrium equations under a symmetrically distributed radial loading (44–46), we obtain the governing equations for *U, V* and *W*:

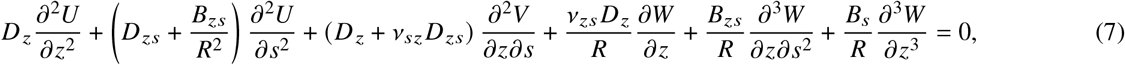

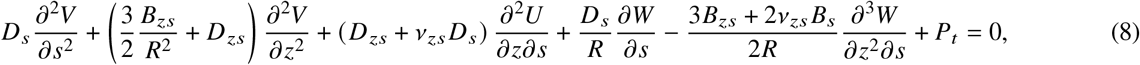

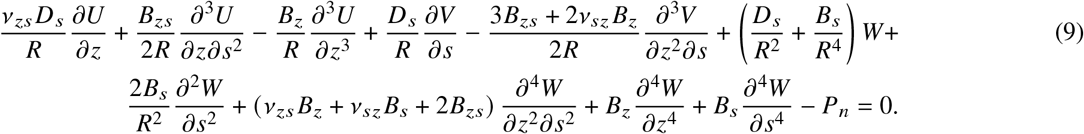

The bending stiffnesses *B*_*a*_ and stretching stiffnesses *D*_*a*_, with *a* ∈ {*z, s*}, are defined as follows:

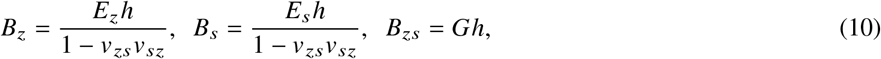

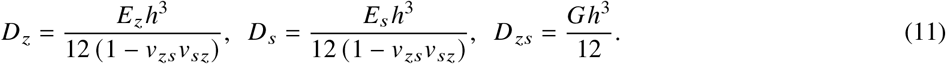

where *E*_*z*_ and *E*_*s*_ are Young’s moduli in *z* and *s* directions, *v*_*zs*_ and *v*_*zs*_ are longitudinal and circumferential Poisson ratios, *G* is the in-plane shear modulus of the cylindrical shell and *h* is the thickness of the shell, and the model assumes the shell thickness is much smaller than the cell radius (*h* ≪ *R*).

To obtain an analytical solution for this problem, the following boundary conditions should be considered:

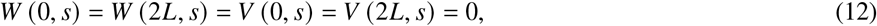

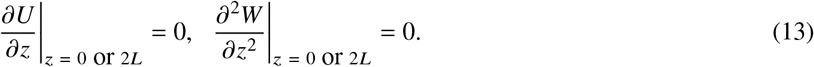

where *L* is the microtubule length.

Taking advantage of the symmetry in the boundary conditions, the single microtubule simulations are performed on only half the shell length. Thus, for convenience, the analytical solution is obtained by considering microtubules of length 2*L*. This linear problem can be solved by utilizing Fourier expansion of displacement variables:

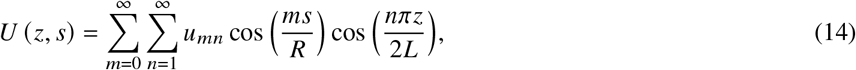

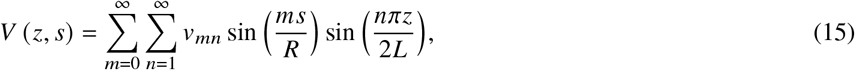

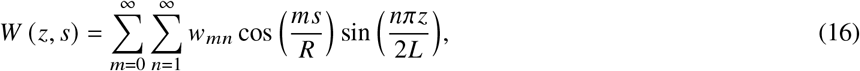

which satisfy the boundary conditions (Eq. 12 and Eq. 13) automatically.

The *s*-dependence of loading *P* dictates that all terms with *m* ≠ 1 must vanish. Focusing on the lowest energy mode *n* = 1, we substituted the resulted expressions into Eqs. 7-9 and obtained a linear system of equations. This system is then solved for *v*_11_ and *v*_11_, taking into account the prescribed geometry, material properties and loading:

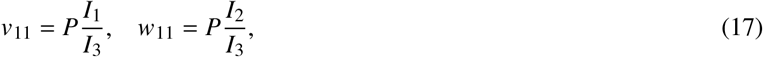

where the expressions for *I*_1_, *I*_2_ and *I*_3_ are provided in § 5.

Maximum lateral displacement of the shell *d*_*theory*_ can be calculated by averaging the sum of displacement components of the central edge in the bending direction over its perimeter:

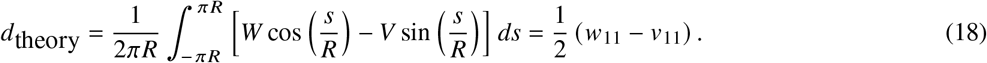

Modeling a microtubule as an Euler-Bernoulli beam with persistence length *p*_*MT*_ undergoing Brownian fluctuations, displacement *d*_*EB*_ is related to the loading as follows:

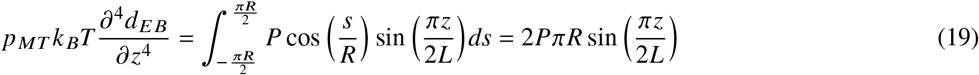

because the load is zero for B outside the internal of *s* ∈[−*ΠR/2, ΠR/2*]. Solving this differential equation by substituting the displacement from Eq. 18, the persistence length of a microtubule is found to be:

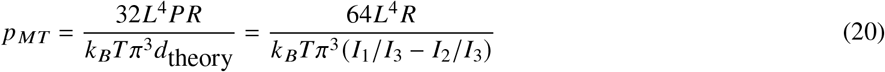

Eq. 20 is fitted to persistence length data of microtubules with varying contour lengths (47) for elastic moduli *E*_*z*_ and *E*_*s*_ as well as shear modulus *G* as fitting parameters (Fig 3a). Thickness *h*, radius *R* and Poisson ratio *v*_*zs*_ were averaged from measurements reported in literature.

## 3 NUMERICAL MODELING

First we validate the applicability of our thin shell theory to cylinders with finite thickness. In these simulations, cylindrical shell of various lengths were subjected to increasing distributed loads P, constrained by Eqs. 12 and 13, and simulated using Abaqus/Standard (48). The resulted lateral deflections were compared to the same analytical setup. The LAMINA material option in Abaqus, which allows for the simulation of orthotropic shell materials, was selected for the analysis.

The cilia microstructure was modeled in Abaqus using serial section electron tomography (SSET) images and data of primary cilia (Fig 2a-c). Microtubule complexes (MtCs) are generated by intersecting two A and B microtubule cylinders (from standard simulations) within *r*_*b*_ center-to-center distance and removing the intersected area of the B compartment (Fig 2f). The decorating axonemal proteins were excluded from our model, and microtubules were tied together at their joint boundaries in our simulations. Nine such MtCs are radially arranged at a distance *r*_*a*_ from the center of the cilium, while being encapsulated by the ciliary membrane (Fig 2e) to form the axoneme. The ciliary membrane is also modeled using the LAMINA material in order to independently enforce the low shear modulus criteria to the model, as implied by the experimental bending modulus *B*_*M*_ and area modulus *K*_*M*_ of the lipid bilayer. The biophysical parameters used for the both the membrane and MtCs are summarized in Table 1.

**Figure 2.**
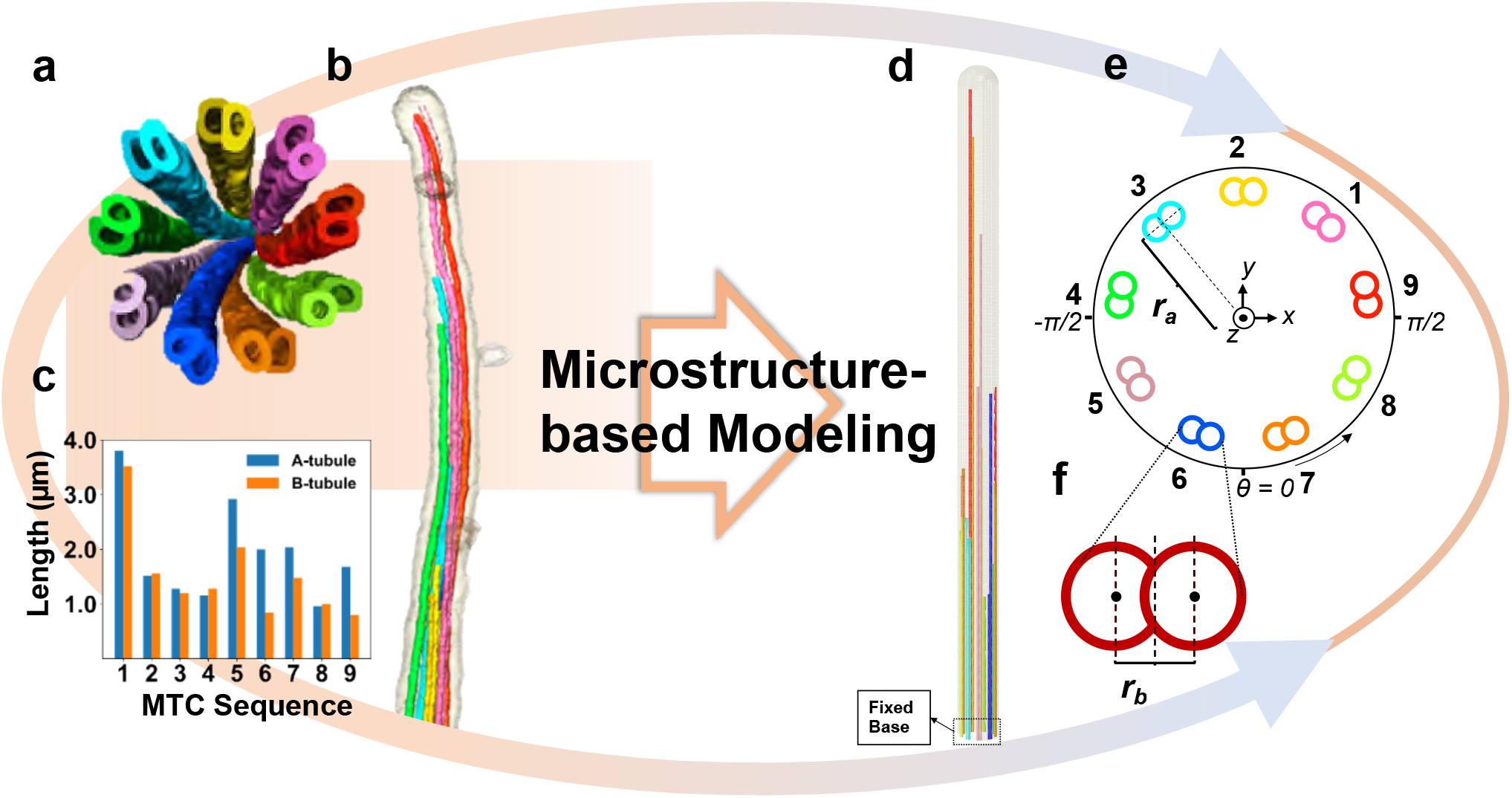
: Serial section electron tomography (SSET) images of primary cilium are provided in cross-sectional (a) and full body (b) views. In addition to providing insights into the topological arrangements and structures, SSET imaging enabled the researchers to quantify the length profile of all nine microtubule complexes (c). (d): Utilizing this information, microstructure-based modeling was employed to develop computational models that consider the interaction between the ciliary membrane and axoneme as separate components. (e): Microtubule doublets are arranged radially, with a distance of 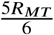 from center. The origin of circumferential direction, *θ* = 0 is designated with respect to cartesian coordinates. (f): Each complex is formed by intersection of two microtubules, with a center-to-center distance of 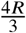.

**Figure 3.**
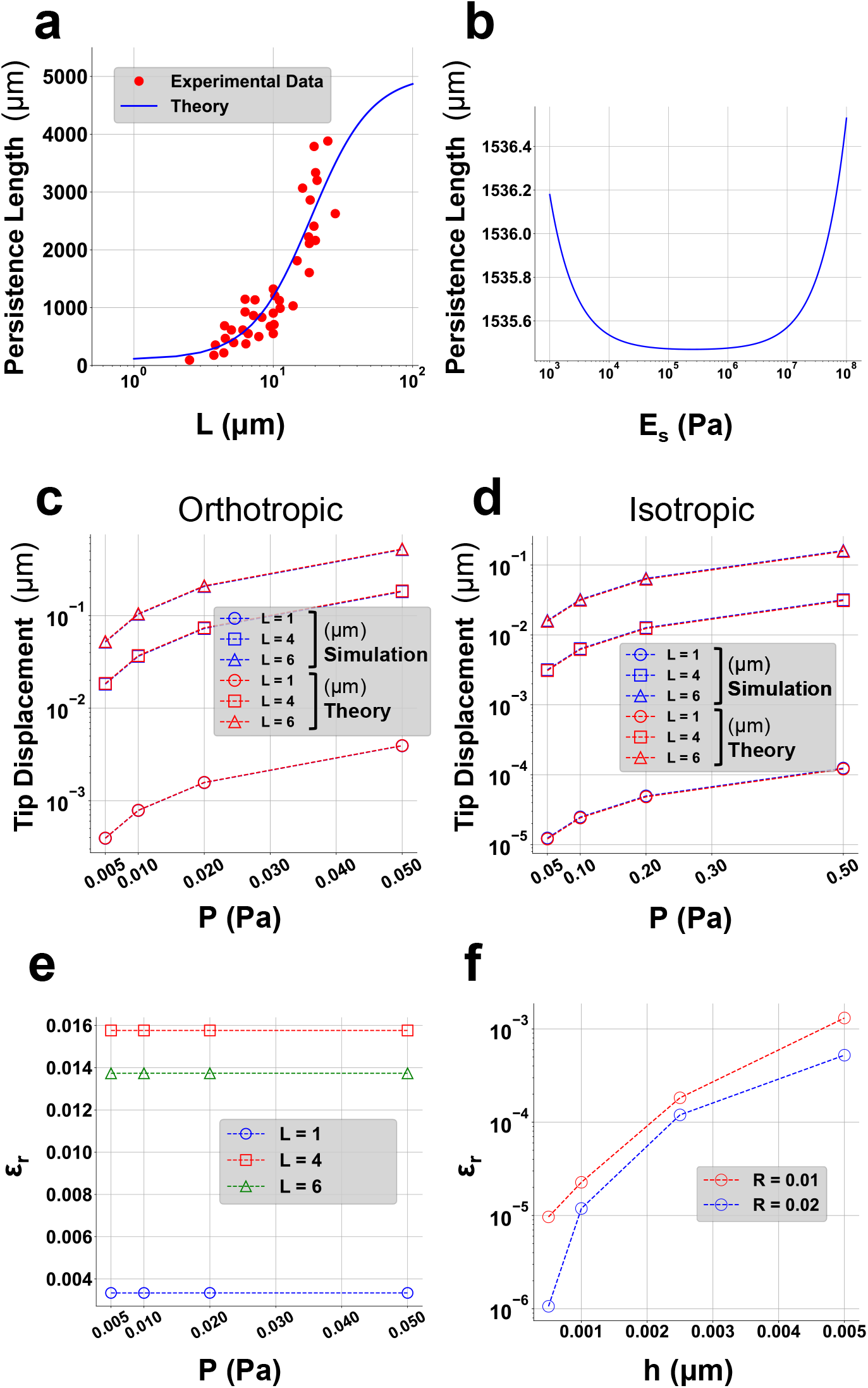
: Comparison or analytical and numerical results. (a) Experimental measurements of microtubules with different contour lengths (47) are used to fit the persistence length from Eq (13) and estimate the elastic and shear moduli: *E*_*x*_ = 2.6×10^9^ Pa, *G* = 7×10^3^ Pa. (b): Due to the relatively small values of h/R and R/L ratios in microtubule shells, varying *E*_*s*_ within its biological range does not significantly alter the persistence length. Therefore, an average value of *E*_*s*_= 2 MPa is used for simulations. (c) and (d): Tip displacements of the microtubules in theory, *d*_*theory*_ and simulation, *d*_*stimulation*_, are shown for fitted (orthotropic) elastic moduli (c) and isotropic inputs (d) under varying lengths *L* ∈ 1, 4, 6*μm* and increasing distributed load *P* ∈ 0.005, 0.01, 0.02, 0.05 Pa. For the isotropic case, *E*_*z*_ = *E*_*s*_ = 2.6*G* and *G* = *E*_*z*_/2 (1 +*v*_*zs*_). (e): For all isotropic simulations, the relative error remained at 3%. However, for orthotropic cases, the relative error increased with length, but it was still less than 2%. (f): The relative error also decreased as the *h*/*R* ratio in the simulated shells decreased, as the theory assumes a negligible thickness

Two cases were investigated in this paper. In the first case, referred to as the representative case, all microtubules in a cilium were assumed to have the same length, which was set equal to the length of the cilium itself. For the second case, we utilized the SSET microtubule length profiles to investigate the effects of microtubule length distribution on overall ciliary stiffness and phenomena associated with axoneme-membrane contact. Fig 2d shows a finite-element primary cilia microstructure model built in Abaqus using data in Table 1 and length profile information in Fig 1c.

The base of the cilium (membrane and axoneme) is fixed in all directions. A spherical cap is positioned on the head of the cilium structure to accurately match Cryo-electron microscopy (EM) images and serve as a vessel for simulating optical tweezers tip displacement experiments. Initially, when the axoneme is upright, a surface traction equal to 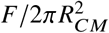 is applied on the surface of the cap in the Cartesian G-direction (Fig 2e) where F represents the total force. As cilium bends, the force direction is allowed to rotate along with the bent cilium in order to prevent stretching of the body.

## 4 RESULTS

We first compared our analytical solutions with finite element simulation of a microtubule singlet as an orthotropical cylindrical shell in § 4.1. We obtained good agreement between finite element simulation and the analytical solutions for this test. We then simulated the entire cilium as nine microtubule doublets and surrounding cilium membrane, and considered their large-deformation and contact interactions in § 4.2. We calculated the effective bending stiffness and persistence length, and explored their length and deformation dependencies in § 4.3. We also investigated the deformation modes of microtubules in different force regimes. Finally in § 4.4, we examined the stress condition within the cilium membrane, especially the maximum stress location and its potential implications for mechanosensing.

### 4.1 Comparison between analytical solutions and simulations of bending microtubule singlets

Fig. 3a shows the persistence length *p*_*MT*_ in Eq. 20 fitted to experimental measurements of microtubules with different contour lengths *L. E*_*z*_ scales the *p*_*MT*_ vs *L* curve and is constrained by the loose maximum plateau of 5000 *μm* measured from experiments. On the other hand, Eq. 20 shows *E*_*s*_ does not significantly impact the persistence length of the cylindrical shell within its measured values in experiment because of the large *L* to *R* ratio considered in our cases (Fig. 3b). *G* is fitted to Eq. 20 using least squares analysis. The resulting material and geometric parameters are summarized in Table 1. To test the comprehensiveness of the theory, two cases were simulated. In the first case, the microtubule properties were assumed to be orthotropic with estimated moduli. In Fig. 3c, the tip displacement of the simulated microtubules for this specific case is depicted alongside the same setup from in Eq. 18 where microtubules of different lengths are subjected to an increasing distributed load P. In the second case, the microtubule was assumed to be isotropic. The fitted longitudinal elastic modulus *E*_*z*_ was used as the Young’s modulus, and the shear modulus was calculated as 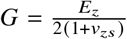. Fig. 3d depicts the same analysis as Fig. 3c, but for the isotropic case. The relative error between the tip displacements obtained from theory and simulation 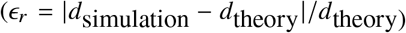 remains approximately 3 percent for all lengths in the isotropic case. However, for orthotropic inputs, this value varies with the length of the microtubules, as shown in Fig. 3e. The discrepancy arises from the fact that the theory assumes an infinitesimally thin shell, while in the simulations, we considered the actual microtubule thickness, denoted as *h*. Nevertheless, the relative error *∈*_r_for microtubules involved in primary cilia simulations remains below 2 percent, which we considered to be acceptable. Fig. 3f illustrates the decrease in relative error with the ratio of microtubule thickness to radius in the simulation. As the theory assumes an infinitesimally small thickness for the shell, when we decrease the thickness of the shell in our simulation while keeping the setup fixed, the resulting displacement approaches the theoretical prediction.

### 4.2 Deformation of the whole cilium and its components

After the careful validation of our finite element solution by the orthotropic cylindrical shell theory, we assembled nine MtCs and put them inside an axonemal membrane to construct a whole cilium model. By leveraging the elastic and shear properties derived from the theory for microtubules, we can represent the MtC assemblies within cilia in their ultra-structural form.

The primary cilia were modeled to include the ciliary lipid bilayer membrane and nine microtubule complex structures. In our model, we considered the role of the transition zone, which anchors the microtubules to the membrane at the base of the cilium and helps prevent lipid diffusion from the ciliary body into the cell membrane. While we excluded the direct contribution of transmembrane proteins in our model, we acknowledged their potential impact on the lipid membrane by considering the combined effect of both transmembrane proteins and the transition zone in reducing the fluidity of the lipid membrane and enhancing its shear response.

The primary cilia of various lengths, along with different microtubule length profiles, were subjected to bending by applying varying surface traction on the spherical cap head. These simulations allowed us to explore the interaction between each microtubule complex and the ciliary membrane during deformation, providing insights into potential phenomena that may have biological implications. By studying these interactions, we aimed to gain a deeper understanding of the mechanical behavior of primary cilia and its significance in mechanosensing processes.

Upon initial observation, the bending of cilia can be classified into two distinct deformation regimes based on the involvement of microtubule complexes. In the first regime, only some of the microtubule complexes contribute to the deformation, while in the second regime, all microtubule complexes actively participate in the bending process. This differentiation allows us to analyze and understand the mechanical behavior of cilia in a more comprehensive manner.

Fig. 4a and Fig. 4b depict six cross-sectional slices of a bent representative (all microtubule lengths = cilium length) cilium with a length of 5 um, showcasing both the low and the high deformation regimes. These slices provide visual insights into the structural changes and deformation patterns that occur within the cilia under different bending conditions.

**Figure 4.**
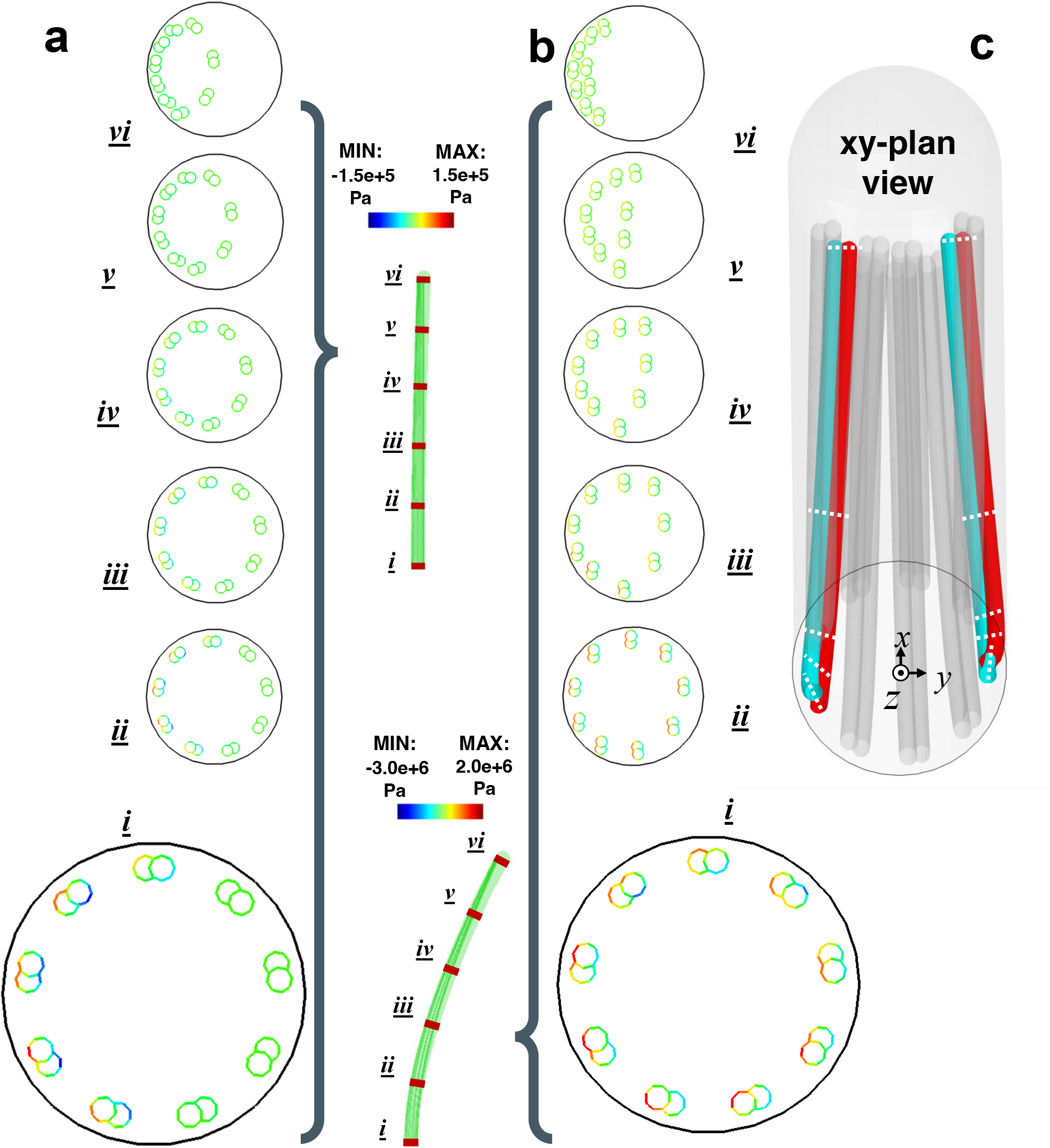
: Deformation of a whole cilium. (a) and (b): The maximum in-plane stress contour is shown for microtubule doublets across six slices of the bent 5*μm* cilium, depicting both low (a) and high (b) bending regimes (one slice magnified to emphasize the differences): In the early stages, microtubule complexes retain their initial orientation while gradually integrating into the mechanics of the cilium. As the cilium bends further, the compressive and tensile stresses exerted on the axoneme base generate a torque that twists the microtubule complexes. This twisting motion maximizes the contact area near the cap of the cilium. Note the large twist that occurs between sections (i) and (ii) in 4(b). (c): Using oblique projection, the twist in two colored microtubule complexes is shown using dashed lines to indicate the rotating doublet axis of symmetry.

In the low deformation case depicted in Fig. 4a, approximately half of the microtubules in the cilia are involved in the bending process, while the remaining microtubules stay undeformed. The axoneme base experiences stress primarily from the side in contact with the bending forces, resulting in tensile and compressive in-plane stresses on the order of 10 kPa.

In the second bending regime, all microtubules in the cilium participate in the bending process, making a more uniform contribution to the overall cilium mechanics. In Fig. 4b, it can be observed that the in-plane stress transmitted to the base of the cilium is approximately 10 times larger compared to the low deformation case. As the MtCs bend, they exhibit a tendency to maximize their contact area with the ciliary membrane, especially when sufficient contact force is applied. This behavior leads to the rotation of the MtCs from their tips, resulting in the generation of a twisting force or torque at the base of the cilium. The rotation of MtCs in the orthogonal direction of the bending can reach up to 90 degrees. Therefore, in addition to the increased in-plane stress, a twisting force is also induced at high deformations. This phenomenon is dependent on the initial orientational arrangement of the MtCs relative to the traction axis. Fig. 4c illustrates the interior of a highly deformed cilium in a 3D view from the bottom *xy*-plane, with two prominently twisted microtubule complexes highlighted with color. It is important to note that the *x*-axis is the bending direction.

### 4.3 Effective cilium bending stiffness

The different low and high bending modes of primary cilia are characterized by their effective bending stiffness *B*_*e*_ and effective persistence length *p*_*e*_. To examine the bending stiffness, we investigated two representative cases with lengths of 4 and 6 *μm*, as well as a 3.85 *μm* cilium constructed based on SSET imaging data. In the simulations we subjected these cilia to an external load of increasing magnitude. The effective bending stiffness of the primary cilia was estimated by treating them as cantilever beams:

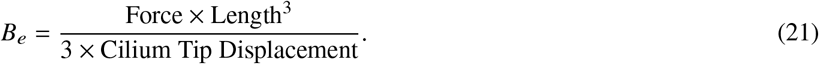

Fig. 5a presents the calculated bending stiffness for the three aforementioned cilia. We observe that, in the representative case where all nine MtCs have lengths equal to the cilium length, the bending stiffness *B*_*e*_ depends on the cilium length. In particular, the representative cilium with a length of 4 *μm*, although comparable in length to the SSET image-based cilium with a length profile for microtubules, exhibits a significantly different bending stiffness. This observation suggests that the overall stiffness of cilia is predominantly governed by the various lengths of MtCs in the axoneme, rather than solely the length of the cilium itself. This observation is further supported by the stiffening of cilia in their low deformation mode in Fig. 5a, where an increasing number of MtCs become involved in the bending process. As more microtubules engage in the deformation, the overall bending stiffness of the cilia increases. Once the axoneme is fully engaged in the bending in high deformation mode, a slight softening of the cilia is observed, which could potentially be attributed to the sliding of the MtCs over the membrane.

**Figure 5.**
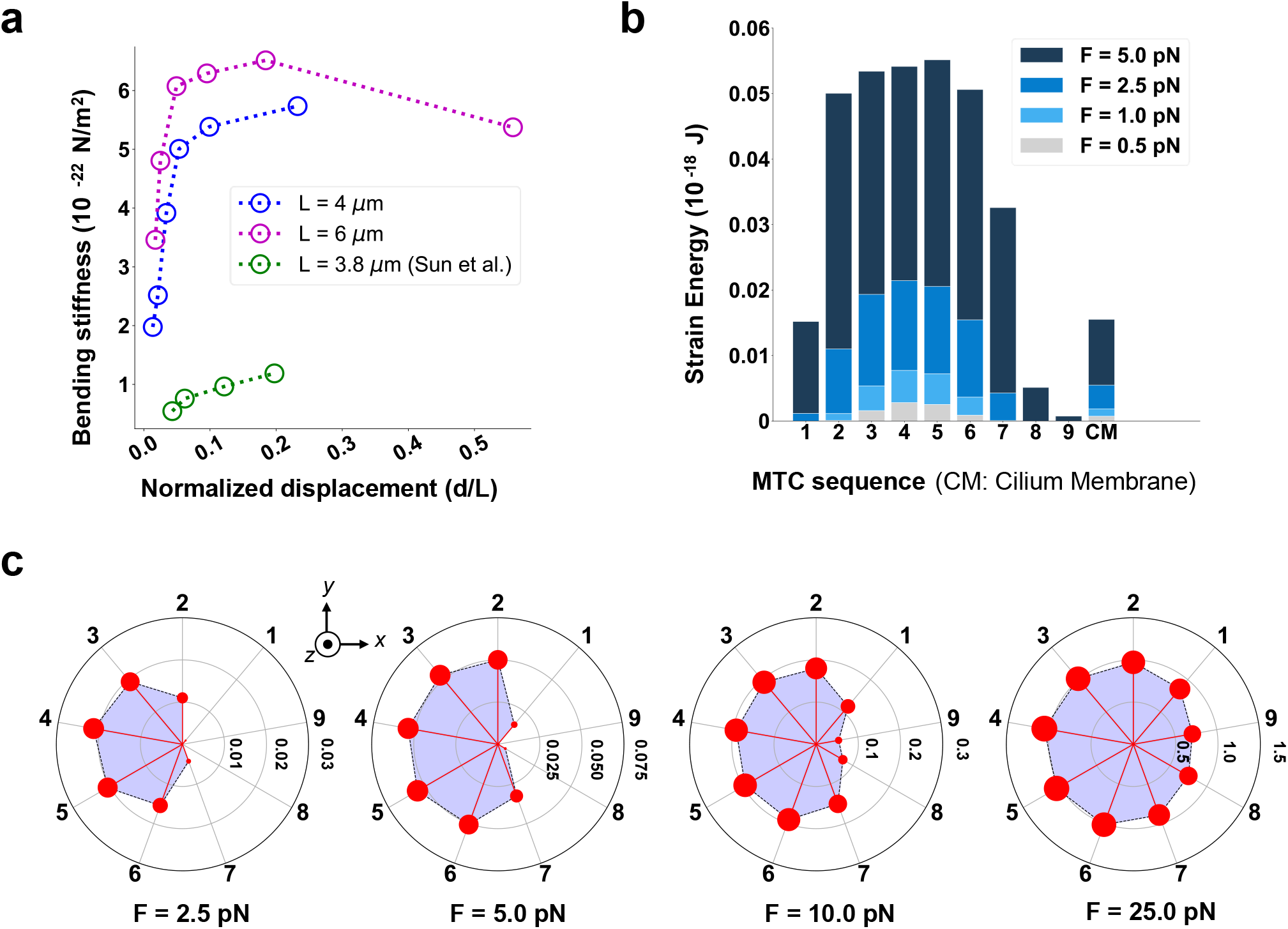
: The length distribution of axonemal microtubules is the primary determining factor of the bending stiffness of primary cilia, rather than just the ciliary length. This is evident from the significant difference in bending stiffness between the representative case and the SSET-based case of comparative length. (a): The increase in bending stiffness with tip displacement indicates the progressive involvement of microtubule complexes in the deformation of the cilium. However, the subsequent decrease in bending stiffness could be attributed to the sliding of microtubules over the ciliary membrane. (b): Microtubules dominate the contribution to the mechanical strain energy, establishing themselves as the primary mechanical components in the system. As a result, they play a pivotal role in bearing the majority of the mechanical load and are instrumental in maintaining the overall structural integrity and functionality of primary cilia. (c): The radial plots depict the gradual increase in strain energy for each microtubule complex as they progressively contribute more to the overall mechanical behavior. This visualization highlights the convergence of strain energy distribution among the microtubule complexes as the cilium bends further.

Fig. 5b provides a visualization of the strain energy increase for each MtC and the membrane in the representative case of a 6 *μm* long cilium under four deformation scenarios. It is clearly observed that the MtCs contribute significantly more to the stiffness of the cilium than the ciliary membrane, highlighting their primary influence in governing the overall mechanics of cilia.

Fig. 5c further illustrates the strain energy distribution in radial plots, allowing for a more detailed analysis of the arrangement-dependent increment of strain energy for MtCs. For the simulation in Fig. 5c a force is applied to the tip of ciliary axoneme from left to right in the positive *x* direction. We observe that the MtCs situated in the left half tend to engage earlier in the bending. By comparing with Fig. 5a, we can clearly observe the strain energy of partially and fully engaged MtCs, which provides further evidence supporting our reasoning for the stiffening and softening regimes.

### 4.4 Stress of cilium membranes and mechanosensing mechanisms

To gain a better understanding of the stress distribution in the ciliary membrane, it is important to quantify both the membrane stress and bending stress within the shell. As showcased in Fig. 6a, the membrane stress is determined by calculating the average in-plane stress across the thickness of the shell. By subtracting the membrane stress from the overall in-plane stress, we can analyze the pure bending stress within the shell. By comparing the membrane and bending stresses, we can gain insights into how the lipid-bilayer responds to bending and the curvature induced by contact with microtubules. This comparison helps us determine the dominant factor in the deformation of the shell when subjected to a certain axial curvature, whether it is the bending stresses or the stretching of the membrane. By analyzing the distribution and magnitude of the bending and membrane stresses, we can assess the relative contributions of these factors to the sensing potential of the ciliary membrane.

**Figure 6.**
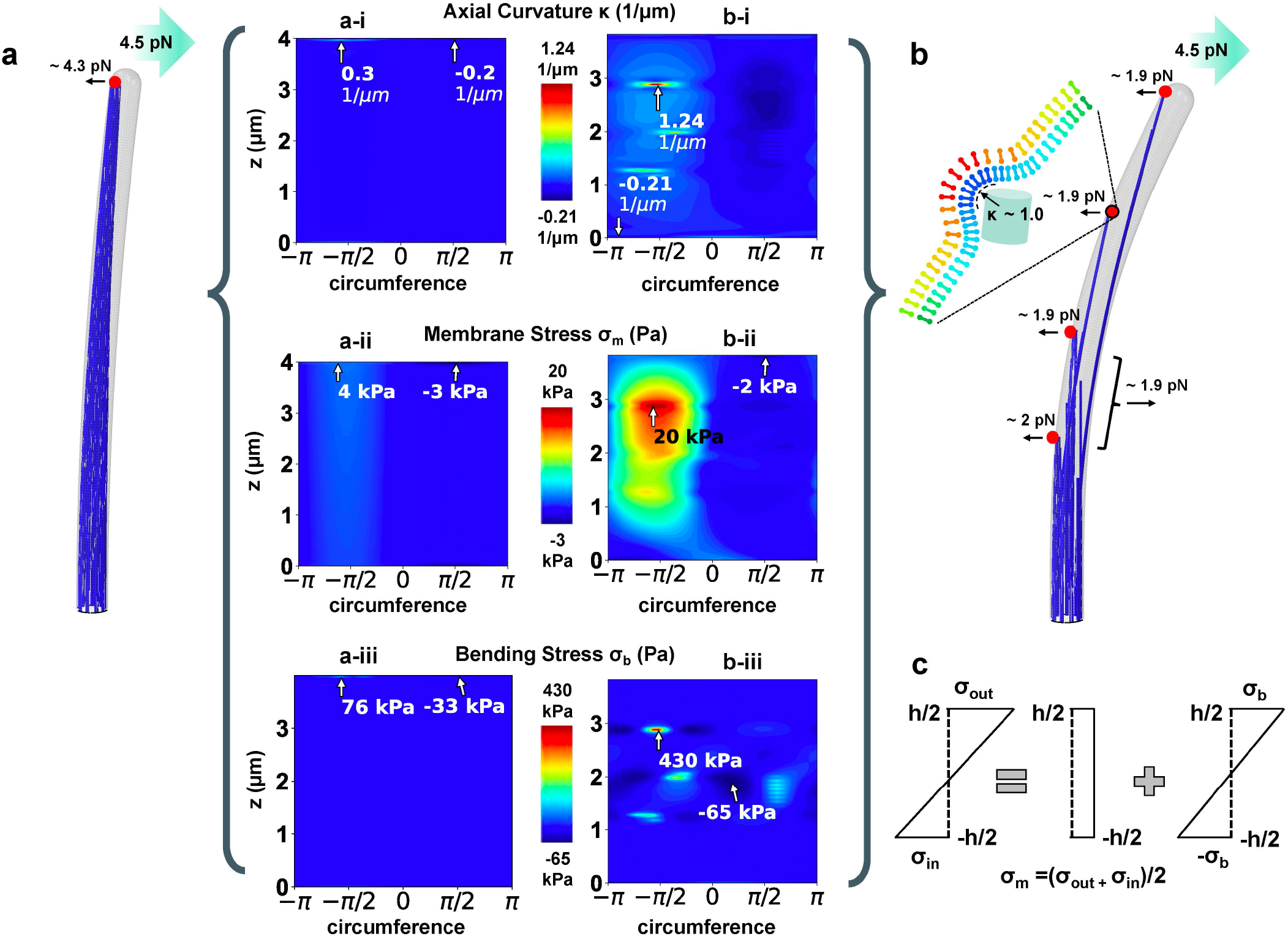
: Distribution of membrane stress, bending stress, and curvature over the ciliary membrane for both the representative case with a length of 4*μm* and the SSET-based case with a length of 3.8 ’m. (a) and (b): While both cases exhibit concentrated stresses and curvature at their respective axoneme-contact focal points, the SSET-based case (b) shows a significantly higher concentration at these points. This is primarily due to the smaller affected area in the SSET-based case compared to the representative case. The maximum curvature and tension points in the SSET-based case suggests a potential mechanism for the activation of piezo channels located on the ciliary membrane. These high mechanical stresses could potentially trigger the opening of piezo channels, allowing for the transduction of mechanical signals into biochemical responses within the cell. (c): Schematic relationship between the total membrane stress as a sum of in-plane and pure bending stresses.

Fig. 6 presents the interaction between the MtCs and the membrane in two different cases: the 4 *μm* representative case and (Fig. 6b) the bent SSET image-based case (Fig. 6c). These interactions are observed under a force of 4.5 pN. In contrast to the representative case, where the contact is mostly localized around the tip of the cilium, the microtubule length profile in SSET-based case exhibits a more uniform distribution of contact force across the membrane. Fig. 6b-i depicts the distribution of axial curvature over the membrane of the representative cilium. As mentioned, the contact in this case is concentrated at the tip of the cilium, resulting in the highest curvature occurring at that location. The curvature peaks observed in the SSET-based cilium are also contact-mediated, but they occur at multiple interaction points. The significant difference in curvature magnitudes between the two cases can be attributed to the smaller contact area in the SSET-based case. This, coupled with greater global curvature induced by the lower bending resistance of this cilium, contributes to a higher overall curvature compared to the representative case.

Although overall the membrane is stretched, as shown in Fig. 6b-ii and Fig. 6c-ii, the bending effect at the same regions is significantly greater than membrane stretching, as shown in the contours in Fig. 6b-iii and Fig. 6c-iii. This observation suggests that the stress gradient within the thickness of the lipid membrane is large, and the shell experiences both tension and compression throughout its cross-section. The distributed contact loads on the membrane generate large curvatures that bend the shell and result in steep stress slope along thickness.

## 5 DISCUSSION AND CONCLUSIONS

We developed an orthotropic shell theory to account for material anisotropy of a microtubule complex and applied the model to a primary cilium. By deriving an expression for the tip displacement of an orthotropic cylindrical shell under a distributed bending load, we were able to formulate an effective persistence length equation. This equation was then used to fit the material properties of the microtubule complex to experimental data on persistence length vs. contour length. Utilizing the obtained material properties of microtubule complex and lipid bilayers, we constructed a microstructure-based model of the primary cilium using SSET images. This model considered the ciliary membrane and axoneme as independent components. Subsequently, we conducted simulations of tip displacement experiments on cilia with varying lengths and subjected to different forces. The goal of these simulations was to investigate the individual contributions of each component and examine the resulting interaction between them.

Our findings show there are strong interactions between microtubule complexes and the ciliary membrane. The microtubule complexes exhibit a tendency to twist upon bending, which increases the contact between microtubules and the membrane. Similar to motile cilia, primary cilia may also possess mechanosensing components. The twisting-induced torque applied to the basal body of primary cilia can initiate conformational changes within the basal body or activate specific pathways associated with it. Hence, in addition to the torque generated by fluid flow over the ciliary membrane, the contact between microtubule doublets and the membrane can also transfer mechanical stress to the basal body.

On the other hand, contact between MtC and membrane gives rise to a high membrane curvature, leading to significant bending throughout the lipid bilayer cross-section around the contact region. This unexpected finding can have profound implications for our understanding of mechanosensing mechanisms. Traditionally, mechanosensitive channels have been primarily associated with membrane stretching or tension, commonly referred to as tension-activated or stretch-activated ion channels. Consequently the location of the largest membrane tension for a primary cilium bent under load is expected to be the most mechanosensitive in the models (49, 50). However, our results demonstrate that the high curvature resulting from local bending of the bilayer can also play a crucial role in activating mechanosensitive channel proteins. This suggests that the channels can be activated by curvature-induced changes in the membrane. This insight challenges the traditional view of ciliary mechanosensation and expands our understanding of the different ways in which cells perceive and respond to mechanical stimuli.

## APPENDICES

Explicit expressions for the parameters *I*_1_, *I*_2_, and *I*_3_ used in Eq. 17 and Eq. 20 are provided here:

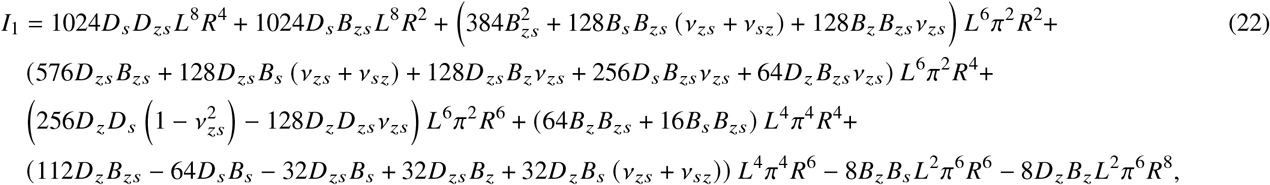

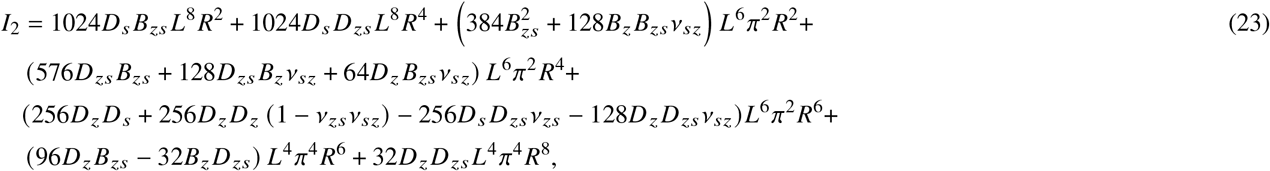

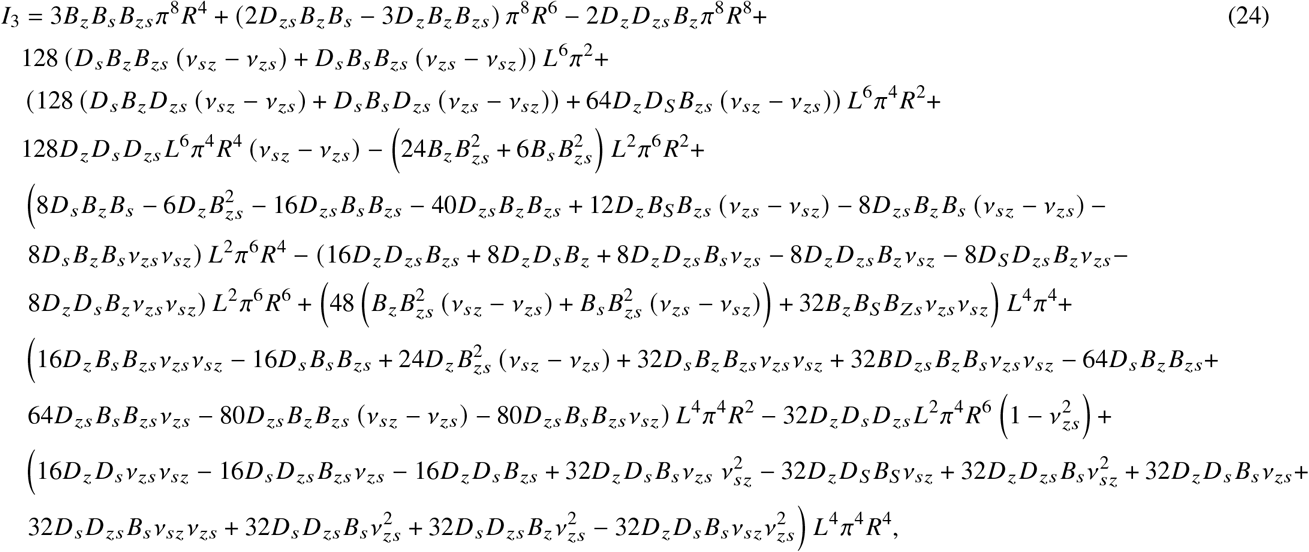

## AUTHOR CONTRIBUTIONS

Z.P., A.R. and Y.N.Y designed the research. N.M. and Z.P. performed the simulations and mathematical analysis, and analyzed the data. All authors contributed equally to writing the article.

## ACKNOWLEDGMENTS

AR was funded by both the National Institutes of Health (NIH 1R15GM132829-01) and National Science Foundation (NSF-DMS 1951568). NM and ZP acknowledge funding from NSF-DMS 1951526. Y-N Y acknowledges funding from NSF-DMS 1951600.

